# Molecular genetic aetiology of general cognitive function is enriched in evolutionarily conserved regions

**DOI:** 10.1101/063636

**Authors:** W. D. Hill, G. Davies, S. E Harris, S. P. Hagenaars, The neuroCHARGE Cognitive Working group, D. C. Liewald, L. Penke, C. R. Gale, Ian Deary

**Author notes:** Authors for The neuroCHARGE Cognitive Working group found in acknowledgements. Correspondence: Dr IJ Deary, Centre for Cognitive Ageing and Cognitive Epidemiology, Department of Psychology, University of Edinburgh, 7 George Square, Edinburgh EH8 9JZ, UK.

## Abstract

Differences in general cognitive function have been shown to be partly heritable and to show genetic correlations with a several psychiatric and physical disease states. However, to date few single nucleotide polymorphisms (SNPs) have demonstrated genome-wide significance, hampering efforts aimed at determining which genetic variants are most important for cognitive function and which regions drive the genetic associations between cognitive function and disease states. Here, we combine multiple large genome-wide association study (GWAS) data sets, from the CHARGE cognitive consortium and UK Biobank, to partition the genome into 52 functional annotations and an additional 10 annotations describing tissue-specific histone marks. Using stratified linkage disequilibrium score regression we show that, in two measures of cognitive function, SNPs associated with cognitive function cluster in regions of the genome that are under evolutionary negative selective pressure. These conserved regions contained ~2.6% of the SNPs from each GWAS but accounted for ~ 40% of the SNP-based heritability. The results suggest that the search for causal variants associated with cognitive function, and those variants that exert a pleiotropic effect between cognitive function and health, will be facilitated by examining these enriched regions.

## Introduction

Individual tests of cognitive function correlate positively, allowing a single latent factor to be extracted from a battery of tests.^1^ This general cognitive factor typically accounts for around 40% of the phenotypic variation in a battery of mental tests and, in large molecular genetic studies, has been shown to be heritable with common genetic variants in total explaining around 30% of phenotypic variation.^2-4^ A higher level of general cognitive function is associated with better health across a range of diseases, both psychiatric and physical, and with lower all-cause mortality.^5^ More recently these phenotypic associations between general cognitive function^6^ and individual tests of cognitive function^7^ with health have been shown to be partly the result of genetic correlations, indicating pleiotropy, meaning that health states show positive correlations with cognitive function in part because the same genetic variants are associated with both cognitive function and health.^6^ However, whereas general cognitive function, along with performance on individual tests of cognitive function, is known to be heritable and to exhibit genetic correlations with health states, for cognitive function, few loci have attained genome-wide statistical significance.^8, 9^ This hampers the effort to understand how genetic variation can result in individual differences in general cognitive function and in turn how this is also associated with variation in health.

This large difference between the variance explained by single nucleotide polymorphisms (SNPs) that do reach genome-wide significance and heritability estimates derived using all tested single nucleotide polymorphisms, indicates that much of the heritability of general cognitive function lies in SNPs that have not attained genome-wide significance. Whereas an increase in sample size will result in an increase in the power to detect significant effects of SNPs in a Genome-Wide Association Study (GWAS),^10^ the problem remains of organising these individual hits into a coherent description of the genetic architecture of cognitive function. Another method that can both increase statistical power and facilitates an understanding of how genetic variation can result in phenotypic variation is Gene-Set Analysis (GSA).^11^ GSA tests the hypothesis that a set of genes, united by a shared biological function^12^ or their previous association with another phenotype,^13^ jointly show an association with the phenotype of interest. The GSA method exploits phenotypes where a highly polygenic architecture is evident by summing the small effects of multiple variants located within the predefined gene set. GSA is not reliant on any single variant attaining genome-wide significance. As such, statistical power is increased as the number of statistical tests is reduced and individual weak effects are combined together to produce a stronger association signal.^13, 14^

Multiple methods exist for the analysis of groups of SNPs treated as the unit of association.^11, 15^ However, many methods assume only a single causal SNP in each of the genetic loci^16^ which, along with being more likely to inflate the type 1 error rate,^17^ also fails to model a polygenic architecture. Other methods suffer from limitations such as requiring access to participants’ genotypes,^18^ and other methods fail to account for linkage disequilibrium (LD) that can lead to the same SNP being counted multiple times within a single gene set.^19^ Here, we make use of a recently-developed method, stratified linkage disequilibrium score regression,^20^ that requires access to only genome-wide association summary level data. This method utilises information from each SNP in a functional category whilst explicitly modelling LD to show if a category is associated with a greater proportion of the heritability of a trait than the proportion of SNPs it contains would suggest. We apply this method to the current largest GWAS on general cognitive function.^8^ We also examine specific tests of cognitive function using the UK Biobank data set. We search for an enrichment in the heritability found in 52 regions corresponding to functional annotations from across cell types, and 10 corresponding to cell-specific functional groups (see Supplementary Table 1). We examine these functional categories, because the distribution of significantly associated SNPs from hundreds of GWAS have indicated that, across diverse phenotypes, significant associations are more likely to be found in regulatory regions such as DNaseI hypersensitivity sites (DHSs)^21^ as well as in protein coding regions^22^ and untranslated regions (UTRs).^23^ DHSs are regions of chromatin vulnerable to the DNase 1 enzyme, as the chromatin in these regions has lost its condensed structure and leaves the DNA exposed, whereas UTRs are involved in the regulation of translation of RNA. In addition, the role of evolutionarily conserved regions has been shown for disease states and psychiatric disorders,^20^ many of which show genetic correlations with the individual tests of cognitive function used here.^7^ By examining the contributions of each of these functional genetic categories we aim to find regions of the genome that play relatively prominent roles in individual differences in general cognitive function.

## Methods and Materials

### Samples

The data used for this study were the summary GWAS statistics from the CHARGE consortium study on general cognitive function in middle and older age which had a total of 53 949 individuals,^8^ and a study of verbal-numerical reasoning based on the UK Biobank^9^ with 36 035 individuals. We next derived a heritability Z-score for both of the data sets that was defined as the heritability estimate produced by LDS regression divided by its standard error. The magnitude of the heritability Z-score is affected by three properties, sample size, SNP based heritability, and the proportion of causal variants. An increase in these three properties is associated with an increase in heritability Z-score. This indicates that the heritability Z-score is capturing information about the genetic architecture of a trait, with traits that have sufficient power, from sample size, high heritability and a high proportion of causal variants yield the greatest heritability Z-scores.

Here, a heritability Z-score of > 7, as used in Finucane et al. (2015),^20^ was used as evidence for a sufficient polygenic signal within the data set for use with stratified LD regression. The CHARGE data set yielded a heritability Z-score of 10.54 and in the verbal-numerical reasoning test a heritability Z-score of 9.64 was derived. This indicates that both of these data sets can be used with stratified LDS regression.

### Cognitive phenotypes

#### General cognitive function

The CHARGE cognitive working group published a GWAS of general cognitive function in 53 949 middle and older age adults.^8^ General cognitive function describes the statistically-revealed overlap between tests of cognitive function, i.e. people who do well on one type of cognitive test tend to do well on others. The cognitive tests included in the general cognitive components used by the CHARGE cognitive working group’s contributing cohorts generally measure fluid cognitive functions. These are functions assessed by tests that tend to include unfamiliar materials, that do not draw upon a participant’s level of general knowledge, and that tend to show a negative trend with age. Each of the CHARGE GWAS project’s cohorts used a different battery of mental tests. Full details of the tests used to measure general fluid cognitive function in each of the CHARGE consortium’s cohorts can be found in Davies et al. (2015).^8^

#### Verbal-numerical reasoning

The verbal-numerical reasoning tests in UK Biobank consists of a series of thirteen multiple choice questions that are answered in a two minute time period. Six of the items were verbal questions and the remaining seven were numerical. An example of a verbal questions is ‘Bud is to flower as child is to?’ (Possible answers: ‘Grow/Develop/Improve/Adult/Old/Do not know/Prefer not to answer’). An example numerical question is ‘If sixty is more than half of seventy-five, multiply twenty-three by three. If not subtract 15 from eighty-five. Is the answer?’ (Possible answers: ‘68/69/70/71/72/Do not know/Prefer not to answer’). To some extent, these questions draw upon materials and information that the participants should be familiar with and the scores on this test are stable when comparing the means between the ages of 40 and 60 years with a linear decline evident from a comparison between the ages of 60 and 70. Genotype data were available from 36 035 individuals who had completed this test. Full details of the genotyping procedures used for this phenotype can be found in Davies et al. (2016).^9^

### Statistical analysis

#### Genetic correlations between phenotypes

Due to the phenotypic overlap between tests of cognitive function^1^ we first examine the degree to which the VNR measure from UK Biobank overlaps genetically with general cognitive function. Genetic correlations were derived using the summary statistics from general cognitive function in CHARGE and Verbal Numerical Reasoning in UK Biobank sets using LD score regression.^24^ The same data processing pipeline was used here as by Bulik-Sullivan et al.^24^ where a MAF of >0.01 was used and only those SNPs found in the HapMap3 with 1000Genomes EUR with a MAF of >0.5 were retained. The integrated_phase1_v3.20101123 was used for LDS regression. Also, Indels, structural variants and strand-ambiguous SNPs were removed. Genome-wide significant SNPs were removed, as well as SNPs with effect sizes of χ^2^> 80, as the presence of outliers can increase the standard error in a regression model. LD scores and weights for use with the GWAS of European ancestry were downloaded from the Broad Institute (http://www.broadinstitute.org/~bulik/eur_ldscores/).

#### Partitioned heritability

General cognitive function^8^ and verbal-numerical reasoning^9^ were analysed using stratified LD score regression, where we followed the data processing steps of Finucane et al.^20^ Stratified LD score regression belongs to a class of techniques that exploit the correlated nature of SNPs. By performing multiple regressions of GWAS test statistics on stratified LD scores, which describe how well a focal SNP tags other SNPs in the same functional annotation, it is possible to estimate the heritability of a phenotype based on the SNPs within the functional annotation. This heritability estimate is then used to derive an enrichment metric defined as the proportion of heritability captured by the functional annotation over the proportion of SNPs contained within it, (Pr(h^2^)/Pr(SNPs). This ratio describes whether a functional annotation contains a greater or lesser proportion of the heritability than would be predicted by the proportion of SNPs it contains, Pr(h^2^)/Pr(SNPs) = 1. The proportion of the heritability for each category is used as the numerator, rather than the heritability of each category. This is due to Genomic Control (GC) being performed on most GWAS data sets and, as a result, the attenuation of the heritability estimate affects the total heritability and the heritability of each SNP set equally. As these are biased in the same direction and by the same amount, the proportion of heritability accounted for by each SNP set remains unaffected by the GC correction although the absolute heritability may change. Stratified LD Scores were calculated from the European-ancestry samples in the 1000 Genomes project (1000G) and only included the HapMap 3 SNPs with a minor allele frequency (MAF) of >0.05.

The same functional annotations as those reported in Finucane et el.,^20^ were used. Firstly, SNPs were assigned to a set of 24 overlapping publically available functional annotations. Supplementary Table 1 details the full set of these functional categories, as well as the references used to construct them. An additional 500bp window was placed around these annotations in order to prevent estimates being biased upwards by capturing enrichment in regions located close to the functional annotations.^25^ A 100bp window was also placed around chromatin immunoprecipitation and sequencing (ChIP-seq) peaks; the inclusion of these additional four sets resulted in a total of 52 overlapping functional SNP annotations which formed our baseline model. In addition, a further 10 sets were examined. These sets consisted of 220 cell-type specific annotations for four histone marks (H3K4me1, H3K4me3, H3K9ac and H3K27ac) which were arranged into 10 broad categories corresponding to histone marks found in the central nervous system (CNS), immune and hematopoietic, adrenal/pancreas, cardiovascular, connective tissue, gastrointestinal, kidney, liver, skeletal muscle, and other. The SNP sets examined here are not independent and the same SNP can appear in many of the sets examined here. The size of each of the SNP sets can be found in Supplementary Tables 2 and 3.

The 10 broad cell-type categories were then analysed by adding each of them to the full baseline model. This resulted in 10 additional tests which included the baseline model and one of the 10 cell-type specific groupings. In this way enrichment for these cell-specific annotations was not driven by their also being a part of the baseline model. Multiple testing was controlled for by using FDR correction applied to these 10 tests.

## Results

### Genetic correlation

LDS regression was first used to determine the degree of overlap between the two cognitive phenotypes used. The genetic correlation between general cognitive function and Verbal Numerical Reasoning was r_g_ = 0.783, SE = 0.056, *P* = 4.63 × 10^−45^ indicating that many of the same genetic variants are involved in both of these traits.

### Partitioned heritability

#### General cognitive function in the CHARGE consortium

Significant enrichment was found for 10 of the 52 functional annotations (Supplementary Figure 1 and Supplementary Table 2). Consistent with many quantitative traits,^20^ SNPs that are found in evolutionarily conserved regions showed a high level of enrichment, where 2.5% of the SNPs accounted for 49.2% of the heritability yielding an enrichment metric (defined as Pr(h^2^)/Pr(SNPs) of 18.87 (SE = 3.91), *P* = 4.88×10^−6^. Statistically significant enrichment was also found after a 500 bp boundary was set around these regions.

Enrichment was also found for two of the histone marks, H3K9ac, where 46.3% of the heritability was found for 12.6% of SNPs (enrichment metric = 3.68, SE = 1.01, *P* = 0.008), and within 500bp of H3K4me1, where 60.9% of the SNPs collectively explained 87.5% of the total heritability (enrichment metric = 1.44, SE = 0.15, *P* = 0.004). SNPs located within 500 bp of repressed regions showed a significant reduction in the level of heritability they captured. These regions accounted for 71.9% of the SNPs, but only explained 44.3% of the heritability (enrichment metric = 0.62, SE = 0.09, *P* = 2.1×10^−5^).

Statistically significant enrichment was also found for SNPs within 500 bp of weak enhancer regions which comprised 8.9% of the SNPs, that collectively explained 38.1% of the heritability of general cognitive function (enrichment metric = 4.28, SE = 1.03, *P* = 0.001). SNPs within 500bp of the functional category of DNase hypersensitivity sites (DHS) also demonstrated significant enrichment for general cognitive function (enrichment metric = 2.05, SE = 0.33). These regions accounted for 49.9% of the SNPs but captured 100% of the heritability (SE = 16%). Whilst this does appear to be capturing the sum of the heritability present it is not clear if this is biologically meaningful as the inclusion of SNPs within 500bp of this category raises the proportion of SNPs in the category from 17% to 50% indicating that the majority of the SNPs within the larger set are not at DHS. SNPs found within 500 bp of introns were also significantly enriched, accounting for 39% of the SNPs and 56% of the heritability of general cognitive function, *P* = 0.00075.

The results for the cell type enrichment analysis indicated that histones that are marked specifically in cell types of the central nervous system accounted for 14.9% of the SNPs, but 45.0% of the heritability (enrichment metric = 3.03, SE = 0.51, *P* = 6.37×10^−5^). The results for the 10 tissue types can be seen in Supplementary Figure 2. The full results for general cognitive function can be found in Supplementary Table 2.

#### Verbal-numerical reasoning in the UK Biobank sample

The pattern of enrichment followed the same trend for verbal-numerical reasoning (VNR) as for general cognitive function. Four of the five functional annotations found to be significantly enriched in general cognitive function were also enriched for VNR (Supplementary Figure 3 and Supplementary Table 3). For the baseline model, evolutionarily conserved SNPs were found to explain an enriched proportion of the heritability; 2.6% of SNPs were found to explain 41.2% of the heritability (enrichment metric = 15.80, SE = 4.42, *P* = 0.00082). As was found for general cognitive function, SNPs within 500 bp of introns also showed enrichment for heritability (enrichment metric = 1.2, SE = 0.14, *P* = 0.0034). This category contained 39.7% of the SNP that explained 56.2% of the heritability. Unlike general cognitive function, SNPs within 500 bp of the histone mark H3K9ac showed significant enrichment, rather than only SNPs found within this annotation (enrichment metric 2.5, SE = 0.48, *P* = 0.0015). This category contained 23.1% of the SNPs which explained 58.1% of the heritability for verbal-numerical reasoning in the UK Biobank data set.

As was found in general cognitive function, histone marks specifically expressed in the central nervous system were found to contain a greater proportion of heritability (enrichment metric = 3.53, SE = 0.60, *P* = 2.40×10^−5^). The enrichment results for each of the 10 tissue types can be seen in Supplementary Figure 4 and the full results for verbal-numerical reasoning can be found in Supplementary Table 3.

Figure 1 illustrates the significant annotations for general cognitive function and provides a comparison for how well these regions were enriched for verbal-numerical reasoning.

**Figure 1.**
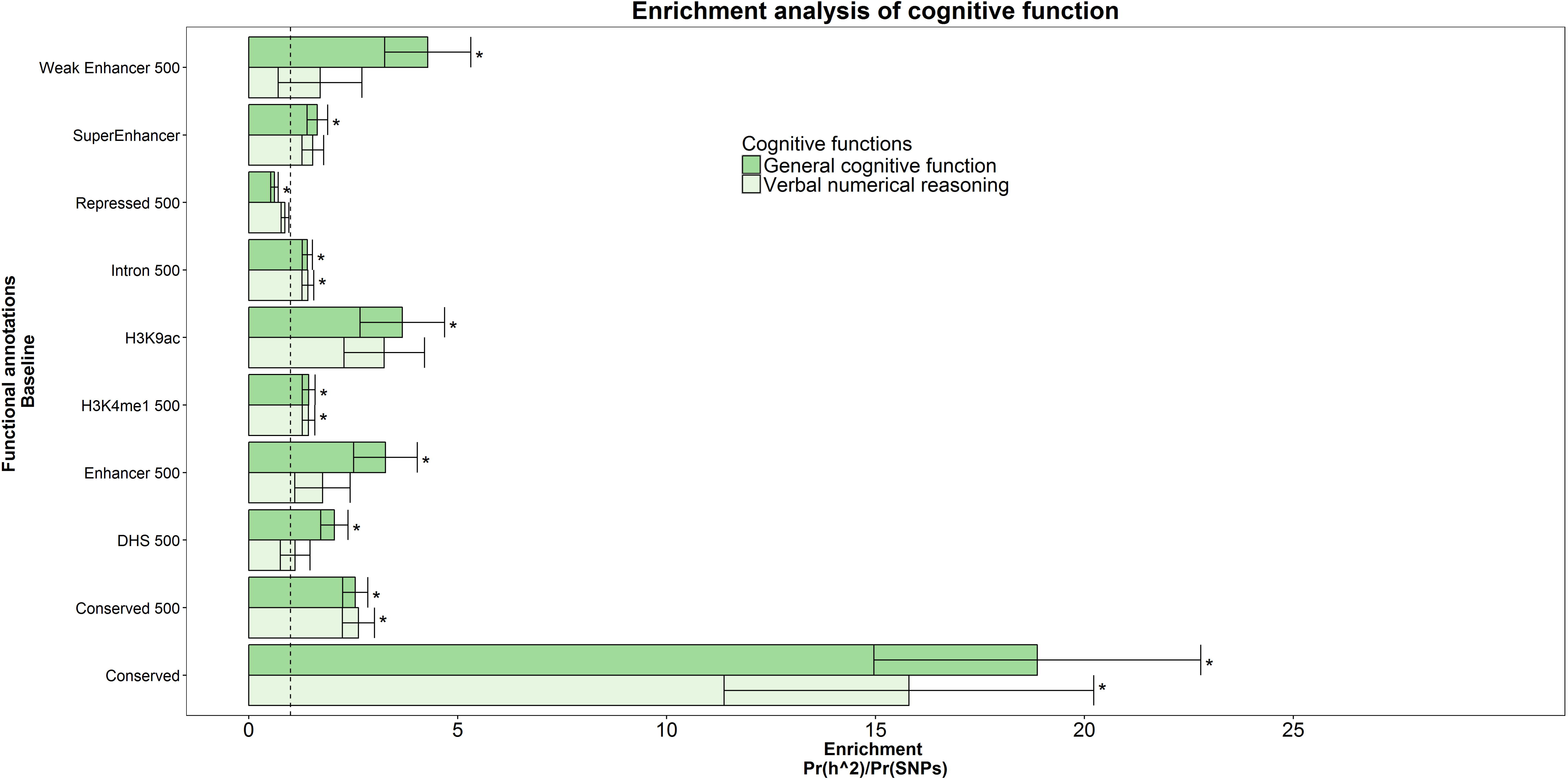
A comparison between the functional annotations that were significantly enriched for general cognitive function. Enrichment was also found in evolutionarily conserved regions for verbal-numerical reasoning. Significant enrichment was also found across the phenotypes for SNPs within 500 bp of introns and within 500 bp of the H3K4me1 histone mark. The enrichment statistic is the proportion of heritability found in each functional group divided by the proportion of SNPS in each group (Pr(h^2^)/Pr(SNPs)). The dashed line indicates no enrichment found when Pr(h^2^)/Pr(SNPs) = 1. Statistical significance is indicated with asterisk.

## Discussion

We partitioned the total heritability found two large, genetically correlated (r_g_ = 0.783, SE = 0.056) GWAS data sets on cognitive function into 24 broad functional annotations and 10 tissue types. Our analysis modelled LD, and took into account overlapping categories, as well as the proportion of SNPs in each category. We make a number of contributions to understanding the genetic architecture of cognitive function.

We find, for both of the cognitive phenotypes examined, the most substantial and statistically significant effects occurred in regions of the genome that are evolutionarily conserved in mammals. The SNPs within these regions accumulate the base-pair substitutions that differentiate species at a lower rate than would be expected under models of neutral selective pressure and these regions are depleted for the number of SNPs compared to regions that are not conserved. This indicates that a large portion of the common variants that are associated with cognitive function are under negative selective pressure. That 40% of the genetic variance in general cognitive function is under negative selection does not imply that higher cognitive function is evolutionarily selected for, but that genetic variance that disrupts the evolutionarily old adaptive design encoded in these regions, thereby decreasing healthy cognitive function, is selected against. This supports the idea that mutation-selection balance plays a substantial role in the genetics of general cognitive function,^26^ particularly when mutational variation is introduced in evolutionarily conserved regions.

Evolutionarily conserved regions are unlikely to be specific to cognitive function, but rather to underlie fundamental design features important for general phenotypic functioning. The evolutionarily conserved regions used in the current paper have also been examined for enrichment with several disease and health related phenotypes. Significant enrichment was found for body mass index, schizophrenia, and HDL cholesterol, but not for coronary artery disease, type 2 diabetes, LDL cholesterol or bipolar disorder.^20^ The diseases and traits that were enriched in these conserved regions each show a genetic correlation with general cognitive function and individual tests of cognitive function (^7, 27^ & Hill et al. in prep), whereas those that showed no enrichment at evolutionarily conserved regions were not genetically correlated with general cognitive function or the verbal-numerical reasoning test used here.^7^ This suggests that not only do these evolutionarily conserved regions of the genome play a greater role in cognitive functions, but they may also harbour variants with pleiotropic effects on cognitive function, health and anthropometric traits, thereby reducing what has been varyingly called system integrity, developmental stability or general evolutionary fitness.^26, 28^

Two previous studies using gene-set analysis have found that gene sets that are conserved between species are enriched for cognitive functions. A study by Hill et al.^12^ found that common SNPs in the *N*-methyl-D-aspartate receptor complex (NMDA-RC) were enriched for general cognitive function in two independent groups. The NMDA-RC is a component within the postsynaptic density (PSD) and using comparative proteomic analysis of the human and mouse PSD it has been found that the molecular composition of the postsynaptic density was highly similar, with more than 70% of the proteins found in the human PSD being found in the mouse PSD^29^ indicating conservation between species. A high level of conservation has also been found between the proteins of the PSD in comparisons between human and chimp (last common ancestor (LCA) ~ 6 million years ago), as well as between mouse and rat (LCA 20 million years ago) and between human and mouse (LCA 90 million years ago), indicating conservation or negative selection indicative of conservation across the mammalian line.^30^

More recently Johnson et al.^31^ used the weighted gene co-expression network analysis to identify a novel module named M3 using cortical brain tissue extracted from living humans during surgery. This module was also present in both disease free humans and in wild-type mouse hippocampi, indicating it had been conserved between both species. In addition, this module was found to be enriched for SNPs associated with general cognitive function and memory in two independent samples. The M3 also mapped poorly onto known biology, including the PSD, differentiating it from the NMDA-RC finding. In the current paper we extend the findings of Hill et al.^12^ and Johnson et al.^31^ by considering all SNPs, not just those found within genes, and show that regions of the genome that are under negative selective pressure harbour an enriched proportion of the heritable variance for cognitive function.

SNPs within 500 bp of introns also showed significant enrichment for general cognitive function and for verbal-numerical reasoning. Enrichment for this region may indicate that, whilst not being translated into proteins, these regions may still exert an influence on individual differences in cognitive function. Indeed, Marioni et al.^32^ have suggested that intronic regions are more likely to harbour genetic variation associated with normal cognitive function than exomic regions. Additionally, SNPs within 500 bp of the H3K4Me1 histone mark were enriched across both cognitive phenotypes. General cognitive function has been shown to correlate highly with tests of crystallised function, and in the present study the genetic correlation with VNR was r_g_ = 0.783 indicating that many of the same SNPs are involved in both facets of cognitive function. This overlap between these two phenotypes may therefore be driven by pleiotropic variants in introns and variants found in the H3K4Me1 histone mark, along with those that are evolutionarily conserved.

The strengths of this study include the use of the largest GWAS of general cognitive function that used established cognitive tests to measure cognitive function. We also use data from the UK Biobank study that includes over 30 000 participants genotyped and processed together using the same VNR test administered in an identical way to in order to remove processing artefacts due to heterogeneity in test used and their administration. In addition the genetic data from UK Biobank were processed in a consistent manner, on the same platform and in the same location.

The limitations of this study include the VNR test used in UK Biobank not being adequately compared to validated psychometric cognitive tests. Also the low response rate in UK Biobank of 5%^33^ indicates that it is not representative of the general population. A further limitation is the use of a general cognitive function phenotype derived from meta-analysis of many smaller studies. Heterogeneity in testing conditions and between different genotyping platforms could introduce a confound that could not have been controlled for here. There may also have been a small number of individuals’ who took part in both CHARGE and UK Biobank. In addition, the stratified LD score regression method is based on an additive model and cannot detect epistatic effects or other sources of non-additive variance. Finally, as with other methods of gene-set analysis, these methods are limited to the availability and accuracy of the annotations used.

Following partitioned heritability analysis we report that regions of the genome under negative selective pressure make a greater contribution to the heritability of cognitive functions than their size would suggest. This indicates that the genetic correlations may not be due to causal alleles being distributed across the genome but, rather, clustering in regions that are conserved. Disease states and anthropometric traits that show genetic correlations with cognitive function also show enrichment in these regions, whereas, in diseases and traits that do not show this pattern of enrichment, no significant genetic correlation with cognitive function is found. Together this suggests that evolutionarily conserved regions harbour variants with pleiotropic effects between cognitive function and as well as diseases and anthropometric traits. In addition, this study aids the search for plausible sets of causal variants by showing that a reduced portion of the genome comprising, only ~2.5 % of the total number of SNPs, can explain around ~40% of the heritability of cognitive function.

## Acknowledgements

Cohorts for Heart and Aging Research in Genomic Epidemiology Consortium (CHARGE) cognitive working group banner includes;

Gail Davies. Centre for Cognitive Ageing and Cognitive Epidemiology, University of Edinburgh, Edinburgh, UK. Department of Psychology, University of Edinburgh, Edinburgh, UK.

Ian J. Deary. Centre for Cognitive Ageing and Cognitive Epidemiology, University of Edinburgh, Edinburgh, UK. Department of Psychology, University of Edinburgh, Edinburgh, UK

Stephanie Debette. Department of Neurology, Boston University School of Medicine, Boston, MA, USA. Institut National de la Santé et de la Recherche Médicale (INSERM), U897, Epidemiology and Biostatistics, University of Bordeaux, Bordeaux, France. Department of Neurology, Bordeaux University Hospital, Bordeaux, France.

Carla, I, Verbaas. Genetic Epidemiology Unit, Department of Epidemiology, Erasmus University Medical Center, Rotterdam, The Netherlands. Department of Neurology, Erasmus University Medical Center, Rotterdam, The Netherlands.

Jan Bressler. Human Genetics Center, School of Public Health, University of Texas Health Science Center at Houston, Houston, TX, USA.

Maaike Schuur. Genetic Epidemiology Unit, Department of Epidemiology, Erasmus University Medical Center, Rotterdam, The Netherlands. Department of Neurology, Erasmus University Medical Center, Rotterdam, The Netherlands.

Albert V Smith. Icelandic Heart Association, Kopavogur, Iceland. Faculty of Medicine, University of Iceland, Reykjavik, Iceland.

Joshua C Bis. Cardiovascular Health Research Unit, Department of Medicine, University of Washington, Seattle, WA, USA.

David A Bennett. Rush Alzheimer’s Disease Center, Rush University Medical Center, Chicago, IL, USA.

M Arfan Ikram. Department of Neurology, Erasmus University Medical Center, Rotterdam, The Netherlands. Department of Epidemiology, Erasmus University Medical Center, Rotterdam, The Netherlands. Netherlands Consortium for Healthy Ageing, Leiden, The Netherlands. Department of Radiology, Erasmus University Medical Center, Rotterdam, The Netherlands.

Lenore J Launer. Laboratory of Epidemiology and Population Sciences, National Institute on Aging, Bethesda, MD, USA.

Annette L Fitzpatrick. Department of Epidemiology, University of Washington, Seattle, WA, USA.

Sudha Seshadri. Department of Neurology, Boston University School of Medicine, Boston, MA, USA. The National Heart Lung and Blood Institute’s Framingham Heart Study, Framingham, MA, USA.

Cornelia M van Duijn. Genetic Epidemiology Unit, Department of Epidemiology, Erasmus University Medical Center, Rotterdam, The Netherlands. Netherlands Consortium for Healthy Ageing, Leiden, The Netherlands.

Thomas H. Mosley Jr. Department of Medicine and Neurology, University of Mississippi Medical Center, Jackson, MS, USA

Correspondence to the neuroCHARGE cognitive working group: Jan Bressler

The work was undertaken in The University of Edinburgh Centre for Cognitive Ageing and Cognitive Epidemiology, part of the cross-council Lifelong Health and Wellbeing initiative (MR/K026992/1). This is supported by funding from the Biotechnology and Biological Sciences Research Council (BBSRC), the Medical Research Council (MRC), and the University of Edinburgh, and supports IJD.

This research has been conducted using the UK Biobank Resource under UK Biobank application 10279. The CHARGE Cognitive GWAS was supported by multiple grants including grants from the NIA (AG033193, AG049505).

WDH is supported by a grant from Age UK (Disconnected Mind Project).

The authors declare no biomedical financial interests or potential conflicts of interest.

UK Biobank received ethical approval from the Research Ethics Committee (REC reference 11/NW/0382).

